# A 3D structural affinity model for multi-epitope *in silico* germinal center simulations

**DOI:** 10.1101/766535

**Authors:** Philippe A. Robert, Michael Meyer-Hermann

## Abstract

Vaccine development for mutating pathogens is challenged by their fast evolution, the complexity of immunodominance, and the heterogeneous immune history of individuals. Mathematical models are critical for predicting successful vaccine conditions or designing potent antibodies. Existing models are limited by their abstract and poorly structural representations of antigen epitopes. Here, we propose a structural lattice-based model for antibody–antigen affinity. An efficient algorithm is given that predicts the best binding structure of an antibody’s amino acid sequence around an antigen with shortened computational time. This structural representation contains key physiological properties, such as affinity jumps and cross-reactivity, and successfully reflects the topology of antigen epitopes, such as pockets and shielded residues. It is suitable for large simulations of affinity maturation. We perform *in silico* immunizations via germinal center simulations and show that our model can explain complex phenomena like recognition of the same epitope by unrelated clones. We show that the use of cocktails of similar epitopes promotes the development of cross-reactive antibodies. This model opens a new avenue for optimizing multivalent vaccines with combined antigen cocktails or sequential immunizations, and to reveal reasons for vaccine success or failure on a structural basis.

## Introduction

Receptor-ligand recognition is at the basis of biological function, and defense by the adaptive immune system. Fast computational tools to undersand evolution or mutation imprint under affinity of binding are needed to understand and exploit the specificity of interactions.

Highly mutating pathogens like HIV, hepatitis C virus or influenza are poorly targeted by vaccines, and elicited antibodies do not necessarily protect again the next mutated strain. Strikingly, a few individuals naturally develop broadly neutralizing antibodies (bnAbs) against a large range of strains [1–5]. Injecting these bnAbs is protective against future infections in certain contexts [6, 7], demonstrating their therapeutic potential.

In view of billions of possible antibody sequences, engineering antibodies with target specificity, desired binding landscape, and no self-reactivity, is a timely and complex challenge. Meeting the requirements for the induction of bnAbs *in vivo*, especially in humans or primates, would support better vaccine design and the development of new bnAbs that are functional but not self-reactive. Antibody responses and ultimately generation of bnAb happens in anatomical structures called germinal centers (GCs), where B-cells selectively mutate their B-cell receptors (BCRs) through somatic hypermutation, which are later secreted as antibodies. High-affinity B-cells are selected for survival and proliferation at the expense of low-affinity B-cells. Consequently, the affinity of the antibodies increases over time, a process called affinity maturation (AM).

The GC’s response to single well-defined antigens has been studied *in vivo* [8]. Many predictive mathematical models have been developed that use abstract antigen–antibody affinities in a probabilistic manner: a mutation is an improvement or decrease in affinity to the single target antigen [9–15]. However, at the scale of multiple or complex antigens, new layers of complexity arise that are not covered by these models. Firstly, mutating pathogens evolve multiple antigenic variants that differ in their accessible epitopes, their frequency, and the amino acid (AA) sequences of their accessible sites. Secondly, during a chronic HIV infection, the epitopes recognized inside the GCs evolve over time and different B-cell clones expand successively towards different epitopes [16]. Thirdly, it has been reported that antibodies produced by GC-derived plasma cells can diffuse back to the GCs themselves and compete for epitope binding of GC B-cells [17], potentially changing the immunodominance landscape of the response [18].

Recent mathematical models have accounted for the quantitative properties of AM towards multiple antigens and compared different vaccine strategies such as simultaneous cocktail immunizations or sequential vaccinations [19]. Each model comes with its own abstract encoding of binding affinities [14, 20–26]. We have previously reported [19] that these representations carry different levels of cross-reactivity. Only some can incorporate differential epitope accessibility, key mutations or shielding, and it is not easy to translate a real antigen or structure into any of them.

A more structural encoding of antibody–antigen affinity would overcome these limitations, and the physiological properties would naturally emerge without the necessity to be manually added to abstract models. Several prediction tools simulate protein folding thermodynamics, sometimes with the knowledge of known antibody–antigen structures [27]. Only a few antibody crystal structures are available so far. Full prediction tools like Rosetta [28], have reached powerful binding accuracy at the expense of few hundreds of computational hours for a single calculation.

One GC typically contains 10,000 cells at its peak. Through mutations and high turnover between proliferation and death, a GC will typically explore 10^5^ to 10^6^ mutations during a single reaction [29]. Therefore, the current available prediction techniques for structural antigen–antibody binding are unable to achieve a single GC simulation in feasible time.

Here, we develop a new hybrid model of antigen–antibody folding that simulates the best folding of the complementary determining region 3 (CDR3) around a predefined antigen structure on a 3D lattice. Real AA sequences are used, up to 12 AAs for the CDR loop, and with complex antigen structures up to 1000 AAs for the antigen. The interaction between neighboring AAs is based on experimentally measured potentials [30]. The hybrid model can derive binding energies in computational times suitable for GC simulations (a few minutes for 1000 affinity calculations). Key properties naturally arise from this representation, such as polyreactivity, cross-reactivity, accessibility and shielding effects, and mutations inducing affinity jumps. As a proof of concept, we adapted and ran GC simulations from an *in silico* model [9] with this structural antigen representation. The model showed physiological GC dynamics and proper AM. We show that the use of cocktails of similar antigens generated favorable conditions for cross-reactivity. GC simulations combined with the 3D affinity model presented here, are suitable for testing vaccine strategies and predicting therapeutic methods to modulate immunodominance in realistic computational time.

We validated the presented method and package on the physiological relevance of somatic hypermutation in GCs, showing it is suited for physiological scale receptor-ligand interactions.

## Results

### A fast computational model for lattice-based antigen–antibody binding

Calculations of antibody-antigen binding energy were so far either simplified to an unphysiological degree or so expensive in computational time that they could not be used in real world problems involving many interactions. Here, we describe a fast algorithm that still captures the important physiological properties.

GC simulations need access to affinity changes by somatic hypermutations. Given a predefined antigen structure (the ‘ligand’), binding energy and affinity have to be computed for a large number of mutated binding regions of the antibody (the ‘receptor’), (Figure 1A). We defined a regular lattice, on which residues can occupy predefined positions on a grid (Figure 1B). Real AA sequences are used, consecutive AAs are on neighboring points and only one AA is allowed on each grid point. A protein structure is represented as a list of moves, namely *Straight, Up, Down, Left, Right* (Figure 1B). All covalent bonds are equal and are neglected. Neighboring but non-covalently linked AAs are assumed to contribute to the binding strength. Their binding energy has been previously estimated from structural databases for each pair of AAs [30]. For two interacting proteins, the binding energy *E*_bind_ is estimated as the energetic sum of individual bonds between the two proteins. The total energy *E*_tot_ of one protein is the energetic sum of interacting and intrinsic bonds, thus, including intrinsic bonds as a stabilization factor of the protein structure (see Methods). We assume that the antigen has a stable conformation, which is not changed by the bound antibody. Instead, the CDR3 loop of the antibody folds onto it. For each CDR3 sequence, a folding has to be generated and the folding structure around the antigen with lowest total energy has to be found (Figure 1C).

**Figure 1:**
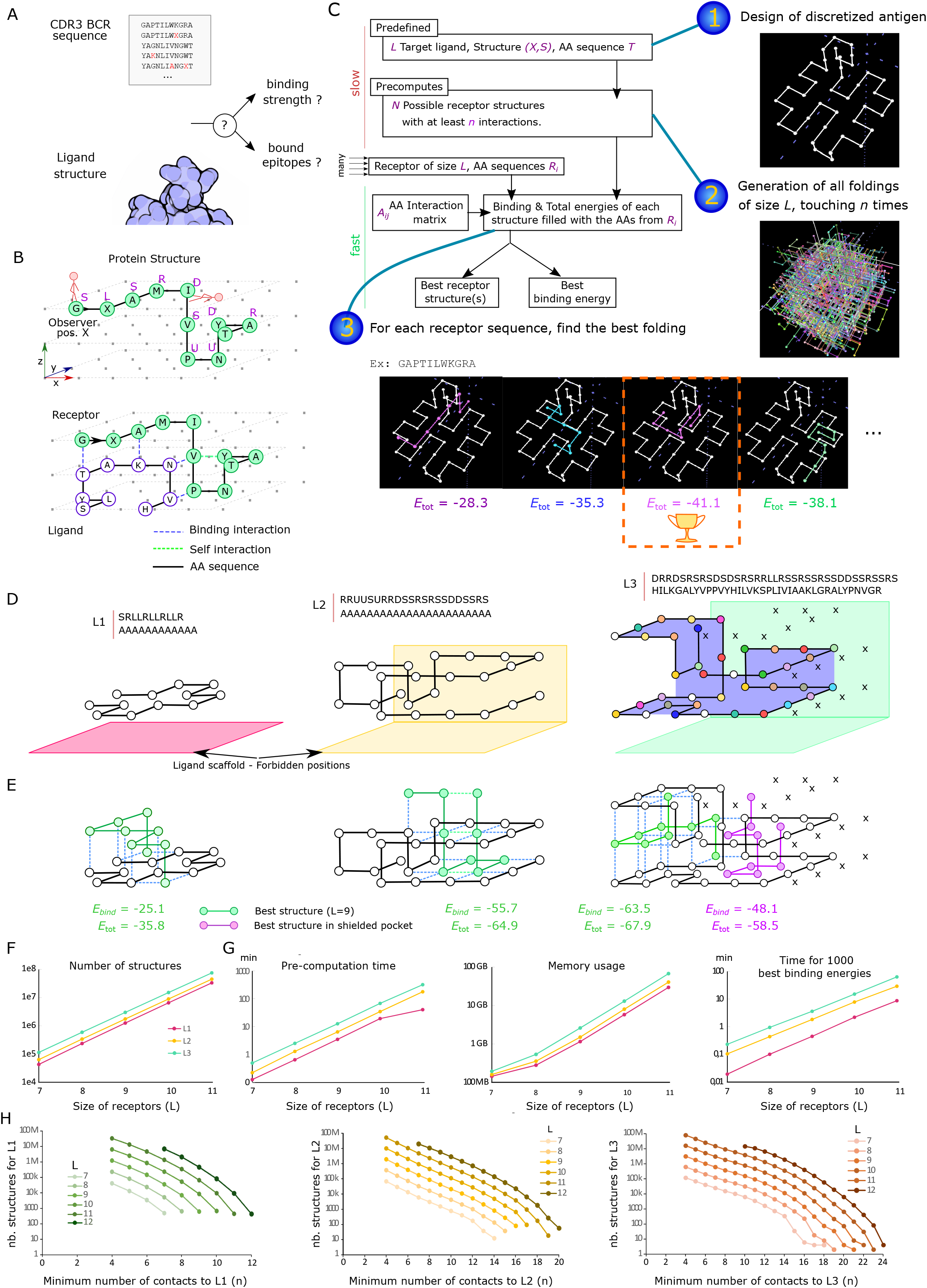
Fast computation of antibody–antigen structural binding on a 3D lattice. **A.** Formulation of the problem: given an antigen, generate a binding landscape of thousands of mutated antibody binding regions, like the CDR3 loop. **B.** Proteins (antigen and antibody binding region) are represented as a path in a 3D lattice, written as: S: *go straight*, L: *turn left*, R: *turn right*, D: *move down* and U: *move up*. The sequence of AAs is shown as a black line. Interactions happen between neighboring AAs that are non-covalently linked, either inside a protein (green) or between the two (blue). The binding energy is the sum of interacting bonds, and the total energy of the receptor additionally includes its intrinsic bonds. **C.** Workflow for the efficient computation of ligand–receptor binding energies. (1) A ligand is predefined with its structure and AAs in the lattice. (2) The set of possible receptor folding structures of predefined length *L*, with a minimum number of contact points *n* to the ligand is pre-computed and stored. (3) For any specified receptor sequence, a binding energy to the ligand *E*_bind_ and total energy *E*_tot_ are derived from each structure of this subset. Then, the ‘best binding energy’ (average binding energy over all structure(s) with minimum total energy) is returned (see Methods). All equally optimal structures are retrieved. **D.** Example of possible ligands: a simple one filled with alanine only (L1), a ligand with a tail and pocket (L2), and a ligand with diverse AAs, a hook, and additional positions blocked to the receptor (shielded residues, ‘X’) that cover the pocket, making it less accessible (L3). Alanine is shown in white; other residues are colored. **E.** For a randomly chosen receptor sequence of length 9, the best folding structures (green for L1, L2; purple for reaching the pocket of L3) are shown with the associated binding energy *E*_bind_ and total energy *E*_tot_ in *kT*. **F.** Number of structures for each ligand, depending on the length of receptors *L* (in AAs), when at least *n* = 4 contacts are required. **G.** Time and memory requirements for pre-computing the structures and then calculating the best energy of 1000 receptor sequences. **H.** Total number of sequences as a function of the minimum requested number of contacts *n*.

Because of the exponentially high number of possible foldings, it is unrealistic to compute all foldings for each CDR3 sequence at every time-step in a dynamic computer simulation of GCs. We developed an optimized algorithm that pre-computes only the interesting possible foldings of CDR3 loops of realistic length *L* between 7 and 14 AAs. Starting from a predefined antigen structure, a recursive algorithm enumerates all possible foldings of the not yet specified CDR3 sequence, with the constraint of touching the antigen at least *n*=4 times (Figure 1C). This considerably reduces the amount of structures to be enumerated and stored in the memory. For each CDR3 sequence to be evaluated, all the previously enumerated foldings are taken one by one and filled with the AA sequence of the CDR3, and the binding and total energies are calculated. As a result, the ‘best binding energy’ is the binding energy of the most stable structure, i.e., the structure with optimal total energy.

As an example, we show three typical antigen structures (Figure 1D): one simple accessible and flat antigen (L1), one antigen with an accessible tail and a hidden pocket (L2), and an antigen with both an accessible hook and a pocket, and additional inaccessible shielded positions, marked X (L3). We assume that positions below the epitope (and on the side of L2 and L3) are inaccessible to receptors, to reflect the antigen scaffold (shown as filled planes). For a randomly chosen CDR3, the best binding structures are shown in Figure 1E, together with the binding energy *E*_bind_ and the total energy *E*_tot_. As expected, the best receptor structure around L2 is located in the accessible pocket. Similarly, the best receptor structure around L3 binds the accessible hook. For comparison, the best structure that would bind the shielded pocket of L3 is shown in purple (Figure 1E) and has much worse binding energy.

For larger CDR3 sequences, up to 75 millions folding structures exist (Figure 1F). The pre-computation time still did not exceed a few hours (Figure 1G) and has to be performed and saved only once. Thousands of CDR3 can be evaluated for best binding energy to these antigens in less than one hour on a single CPU (Figure 1G), implying that it becomes possible to simulate a full GC within a few hours. By increasing *n*, the required minimum number contacts to the antigen, calculations are further speeded up (Figure 1H). For instance, for ligand L3, receptors of length 11 lead to 75 million structures with four contacts, which can be brought down to only 5.6 million by requiring nine contacts. Thus, the structural model for antibody–antigen binding has the capacity to model complex antigens with an efficiency suitable for GC simulations.

### Structural properties of the model

#### Antibody specificity is reflected in the model

Thousands of random receptor sequences of length 7 to 11 AAs were sampled. The best binding energy of the receptors varied according to a broad Gaussian distribution in the range of 20, 40 and 50 *kT* for L1, L2 and L3, respectively (Figure 2A), thus, generating different binding landscapes for each sequence. Despite millions of possible good folding patterns, there are receptors still not matching this antigen. Thus, the calculated best binding energy contains structural information about the receptor sequences. The best binding energy of receptors increased linearly with their length (Figure 2A) because they have more options for binding and a bigger folding ensemble. Therefore, we do not recommend comparing energies between receptors of different lengths, unless a correction coefficient is added.

**Figure 2:**
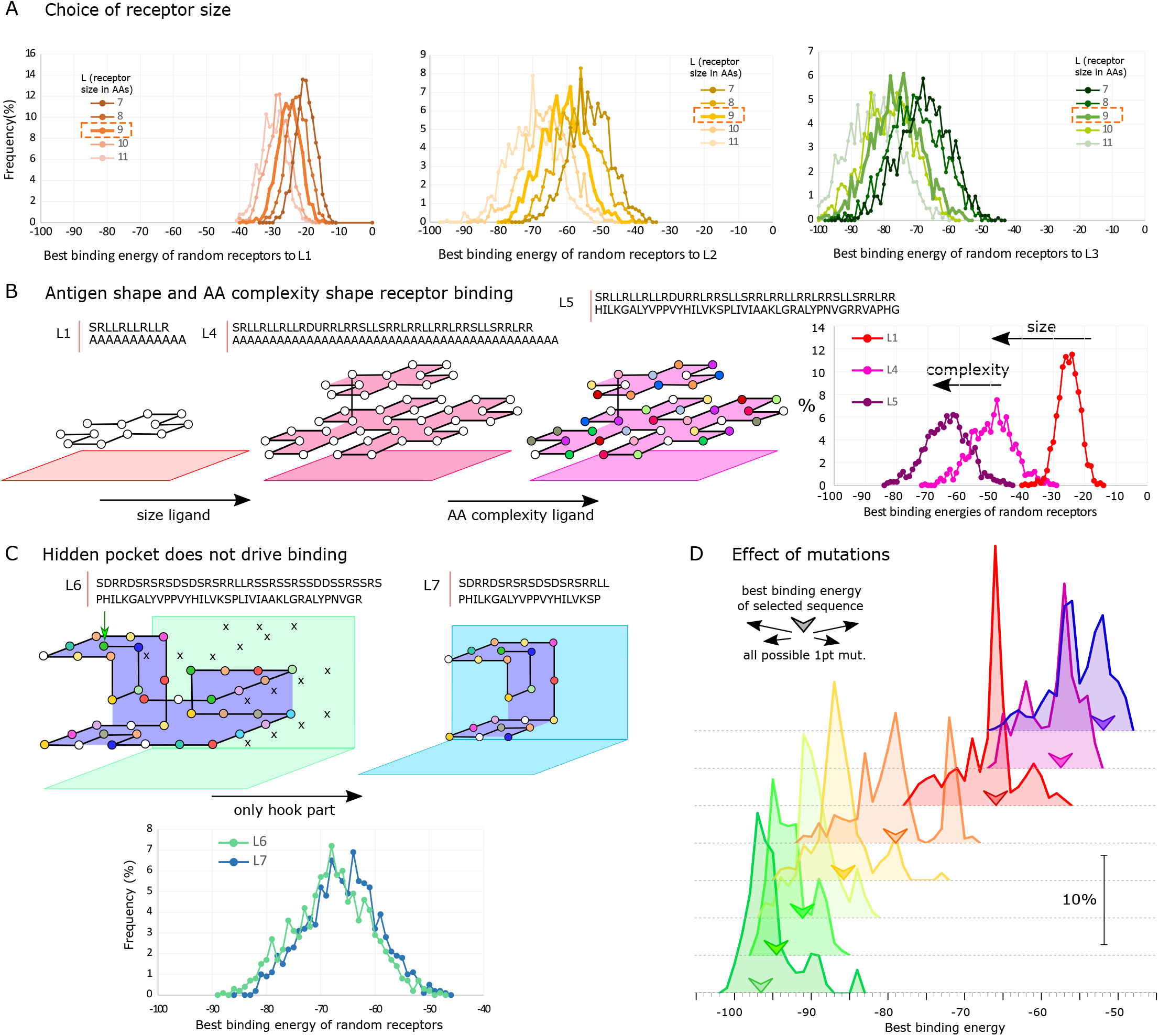
Distributions of receptor binding energies. The best binding energy of 2500 randomly sampled receptors **(A)** of length 7 to 11 to ligands L1, L2 and L3; **(B)** of length 9 to ligands L1, L4 and L5; and **(C)** to L6 and L7. The structure of the ligands L2 and L3 are as depicted in Figure 1D. **D.** Histogram of best binding energies induced by all possible single point mutations, for eight selected receptor sequences with different best binding energies for L6.

#### Steric hindrance of epitope access

The impact of the antigen structure onto the binding energies was investigated on a bigger version of ligand L1 with a similar flat shape. It induced a global increase in binding (Figure 2B), due to more available positions, especially corners where three AAs are accessible from the same point. We therefore suggest comparing ligands of a similar size. In the case of L3, we first created a more physiological variant L6 where the hole in the pocket was filled (green arrow), to avoid receptors going through it. We then monitored the contribution of the shielded pocket by removing it in ligand L7 (Figure 2C). The distribution of best binding energies remained similar because the receptor structures accessing the pocket were not favored. This shows that the model is a suitable tool for studying the effect of steric hindrance of epitopes such as HIV GP120 protein [31].

#### Antigen AA complexity

Changing the AA composition of L1 from alanine only to diverse AAs (L5) (Figure 2B) induced a strong improvement of the best binding energy. If an antigen contains only one type of AAs, receptors do not benefit from different folding structures.

#### Point mutations

In the GC, somatic hypermutation generates point mutations that lead to increased or decreased best binding energies. All receptor sequences for L6 displayed best binding energies between −90 and −45 *kT* (Figure 2C). We tested all possible point mutations for eight sequences of length 9 with different best binding energies to L6 (Figure 2D). Mutations could always do both, increase or decrease the best binding energy, even when starting from sequences with very high (green) or very low (purple) energy. Large energetic shifts of ±15 *kT* were possible by single mutations.

#### GC simulations show affinity jumps and clonal bursts

In order to predict AM and ultimately the efficiency of multivalent vaccines from the structural properties of the antigens, the next step was to simulate full-scale GC reactions. We incorporated the structural binding model into an agent-based model for GC dynamics, where B-cells and T-cells are explicitly modelled to move, interact, be selected, proliferate, die or exit in 3D space [9, 32] (see Methods). The best binding energies derived from the structural binding model were mapped to an affinity (see Methods, Eq. (6)). In the simulation, the affinity between the BCR and the antigen is identical to the probability of capturing the antigen, which is needed later to survive T-cell selection.

With suitable values for these two parameters (see Methods), realistic GC dynamics and AM for a single complex target antigen L6 were found (Figure 3A). The number of B-cells over time in single GCs expands first, reaches a maximum at day 8, followed by slow decay. Starting randomly from cells with at least 0.0001 affinity, the average affinity of B-cells reached 0.3 to 1 after 21 days. The average number of mutations of cells leaving GCs was around six mutations, as estimated in [33]. The high affinity receptor sequences from five GCs were different (Figure 3B). In contrast, in most abstract affinity models the high affinity sequence is unique [19].

**Figure 3:**
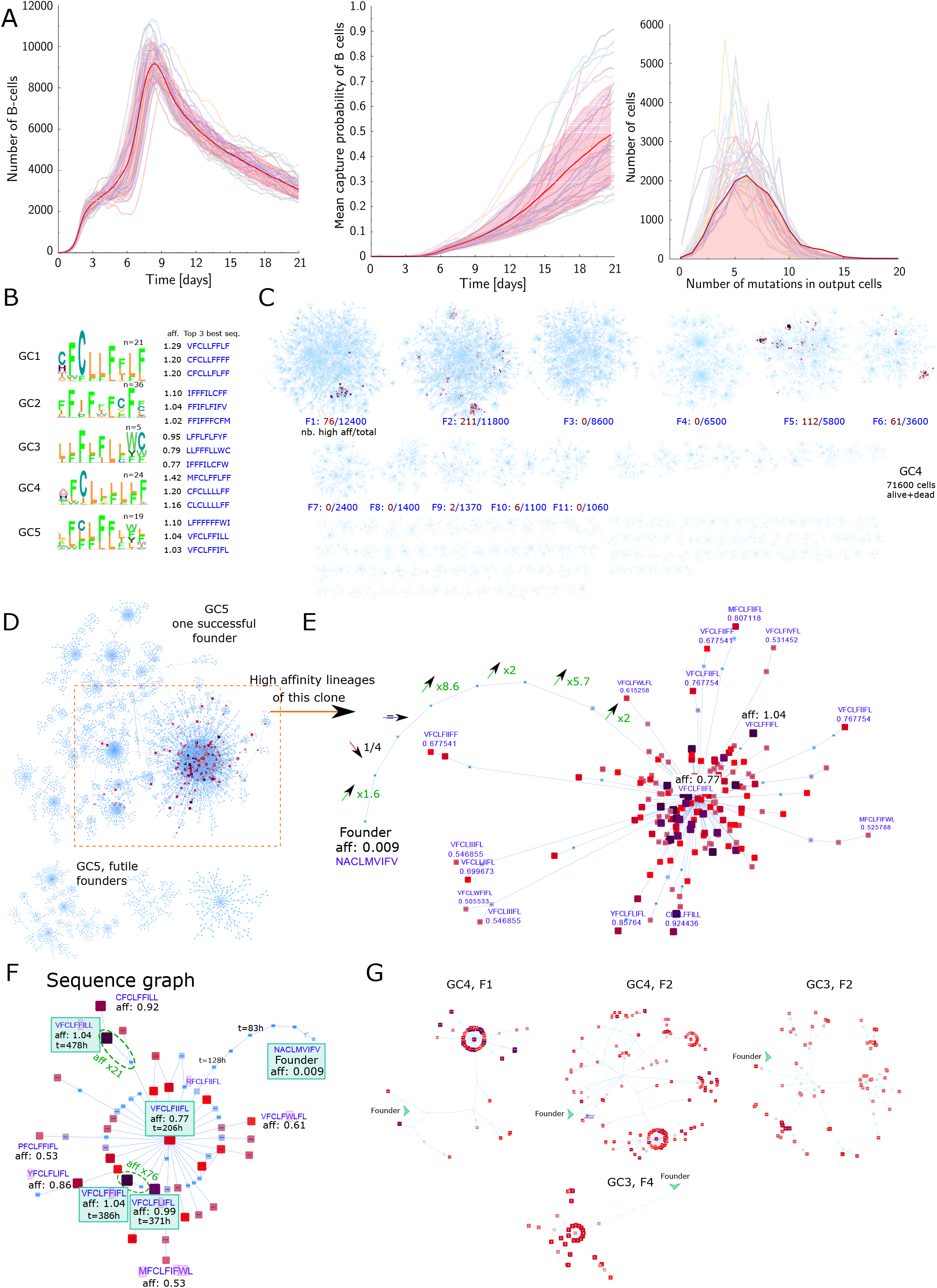
Full scale *in silico* GC simulations and mutation histories of high-affinity cells. **A.** Dynamics of 25 GC simulations against antigen L6, without glycans, and with receptors of size 9, starting from random BCR sequences with at least 0.0001 affinity. The number of GC B-cells (left), the mean affinity of GC B-cells (middle) and the distribution of mutations per GC output cell (right). **B.** The best three sequences and consensus of BCR sequences with very high affinity (> 0.75) from five GC simulations. **C.** Network (forest) representing all the mutation histories of the cells (alive or dead) that existed during a single GC simulation. Nodes are sequences and the edges connect a mutated sequence in daughter cells to the mother sequence. Each panel represents all the progeny sequences of one founder sequence (i.e., one clone). The panels are displayed starting from the founders with most progeny. High-affinity cells (> 0.5) are shown in dark red and are quantified beneath each founder dynasty. **D.** Details of one founder progeny where a big burst could be observed, and the progeny of futile founders that did not reach high affinity. **E.** Lineage history of high-affinity sequences (> 0.5) of the successful founder in D, including the successive affinity increases with each mutation from the founder sequence to the burst. **F.** Same as E, except that identical sequences are merged. The time of first appearance of the sequence and selected large affinity jumps are indicated. **G.** Progeny of high-affinity cells back to the founder cell for different clones in different GC simulations. In all network representations, node size and color gradually represent affinity: light blue for affinities < 0.3, red at 0.6, purple at 0.9, and black for 1.0. The position of nodes is determined automatically by the visualization algorithm Prefuse Force Directed Layout that segregates components of the network in 2D, allowing to see the hubs.

Next, we analyzed all receptor sequences generated by mutations during 21 days of a GC reaction. We retrieved all (around 70, 000) mutations that occurred in selected or dead cells and found that more than one GC founder cell achieved high affinity (Figure 3C), thus, pointing to parallel AM of multiple founder cells *in silico*. This is in accordance with the observation that some GCs are dominated by single clones while others display coexistence of multiple founder clones with high affinity [32].

To characterize the mutation history of high-affinity cells back to the founder cell, we selected the mutation cluster of a winning or losing founder cell in Figure 3D. The former showed an explosive expansion of a sequence into many mutated daughter sequences with high affinity. More bursts (hubs in the trees) happened within both the winning and losing clones (Figure 3C). The history of affinity changes during AM leading to high-affinity sequences revealed that increases and decreases of affinity happened, typically by 2− to 10−fold, before expansion of the high affinity sequence (see founder with affinity 0.009 in Figure 3E). In the simulations, not exclusively advantageous sequences are selected, which well reflects the stochastic nature of selection observed in [32, 34].

In the tree representation, the same sequence can occur in different nodes with a different order of mutations. We simplified the graph by merging identical sequences around the bursting cell (Figure 3F). Strikingly, single point mutations inducing a 74−fold increase in affinity were observed, similar to key mutations like the acquisition of L33 in antibodies against ovalbumin [35]. Thus, the possibility of key mutations is captured by our model.

The comparison of the history of winning founder cells from different GC simulations (Figure 3G) illustrates how different mutation patterns can be. All of them show clonal bursts to rather different degrees. Few reports have studied single cell sequences inside a single GC. Interestingly, these bursts have been observed in a subset of GCs [32] in response to protein antigens. This indicates that our structural model reflects several non-trivial physiological properties of GCs.

#### Antigen cocktails with high similarity promote cross-reactivity

Next, we simulated GCs with two epitope variants with same structure (L6), but more or less sequence similarity (Figure 4A). Interestingly, the GC dynamics and AM were not impacted by the addition of the second epitope variant. For similar epitopes, the simulated GCs could be classified in four groups according to cell affinity to both epitopes at day 11 (Figure 4B). In some GCs, highly cross-reactive cells emerged (GC4), in others the GCs matured to one antigen only (GC2). In most cases, cross-reactive cells were enriched compared with the cells recognizing only one epitope (GC1,3). Therefore, high variability could be observed between GCs and the overall affinity to the best antigen did not reflect the success to generate high cross-reactivity.

**Figure 4:**
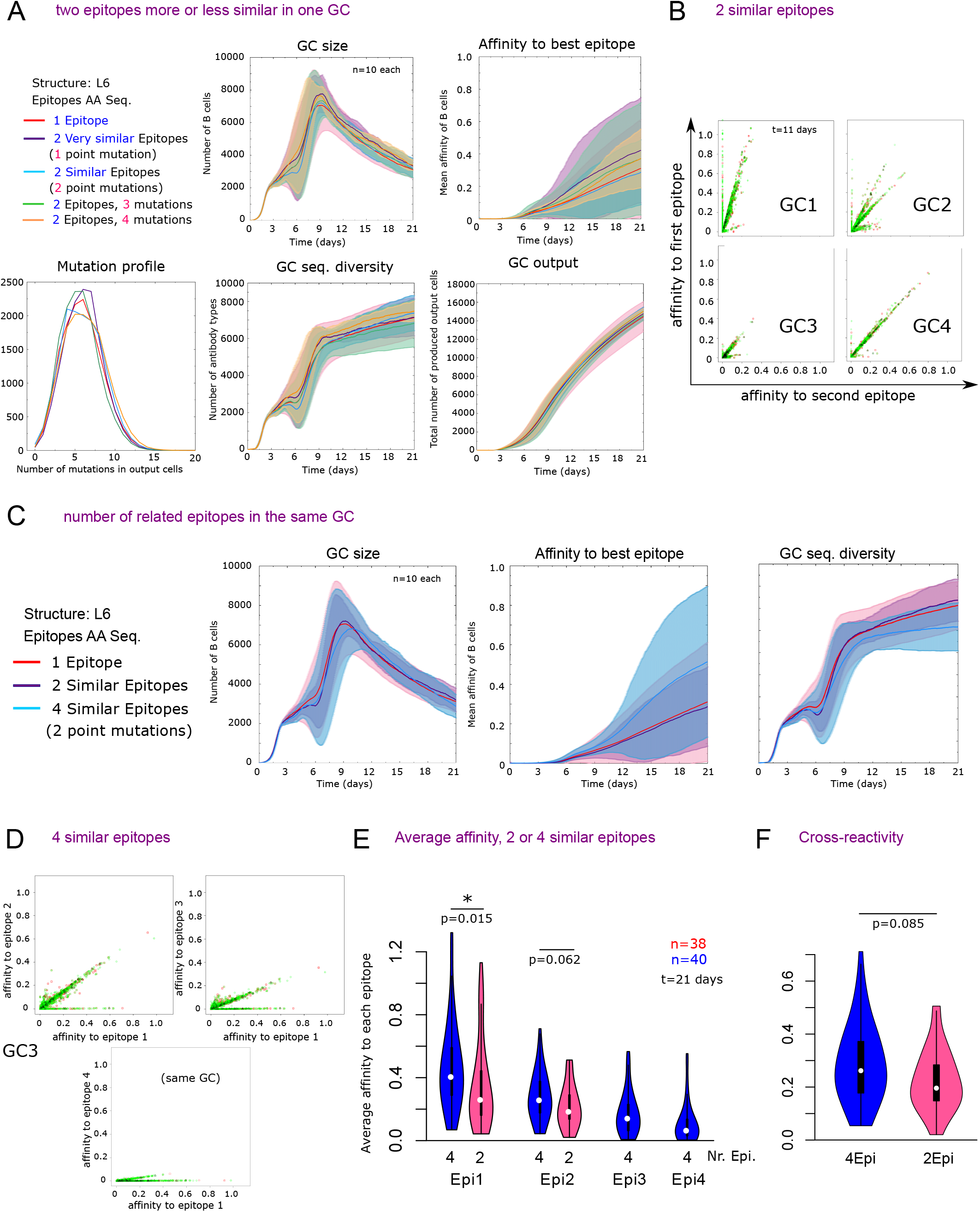
Dynamics and cross-reactivity of GCs with two or four antigens. **A.** Dynamics of GC and affinity maturation for two epitopes variants of the antigen structure L6 that differ by one or two point mutations. **B.** Affinity of single cells in four different GCs imulations to two similar epitopes, which differ by two mutations, at day 11. **C.** GC dynamics with two or four similar epitopes (differing by two mutations). **D.** Affinity of single cells in a single GC to pairs of epitopes at day 11. **E.** Average affinity to each epitope of single GCs with two or four similar epitopes. The epitopes are sorted from highest (Epi1) to lowest (Epi2 or 4) average affinity in each GC. **F.** Average cross-reactivity of single GCs raised with two or four similar epitopes, defined as the minimum affinity to the two best recognized epitopes, for each cell.

Increasing the number of similar epitopes to four did not induce major changes in the GC characteristics (Figure 4C). However, B-cells developed high reactivity to two but not to all antigens (Figure 4D). In order to quantify this effect, we calculated the average B-cell affinity to each epitope for individual GCs with two or four similar epitopes, starting from the best recognized epitope in each GC (Figure 4E). The concomitant use of four similar antigens induced a slightly higher affinity to the two best recognized epitopes as compared to GCs with only two similar antigens. Further, we assessed the average cross-reactivity of each cell for each GC, measured as the minimum affinity to the two best recognized epitopes (Figure 4F). The average cross-reactivity was slightly increased by the combination of four similar epitopes instead of two. Therefore, adding more similar epitopes is not harmful for cross-reactivity but rather supports better recognition of the epitopes.

The model shows that using combination of similar epitopes enhances the chance of generating cross-reactive antibodies, even if less GC founder cell recognizes each epitope.

We believe this will support the design of better vaccines towards poly-reactivity, conserved recognition or broad neutralization.

## Discussion

A realistic representation of antibody–antigen binding that can be simulated efficiently on regular computers was presented. For the first time, an affinity directly derived from a structural *in vivo*-like folding was used in full scale GC simulations of AM. The mutation history and AM can directly be linked to the structural properties of the antibodies–antigen binding *in silico*. This representation can encompass the complex topology of real antigens, and simulate vaccination with either one antigen with complex epitopes or cocktails of many related epitopes. The structural affinity model can be easily transferred to other applications like thymic selection.

Several assumptions were made to achieve efficient computational time. The antigens and receptors were discretized on a 3D grid lattice. Instead of enumerating receptor structures and docking them, we perform both simultaneously, allowing to discard huge amounts of possible structures. We do not pretend to simulate the exact affinity of antigen–antibody interactions: Real affinities will not be captured by simulation of the folding of a single CDR loop around a pre-folded antigen but is influence by many other surrounding factors.

However, critical features and behavior of antibody–antigen recognition are well captured in our system: The binding energy of each individual AA pair is taken from an experimentally derived energy potential [30]. Many non-trivial properties of antibody–antigen binding were successfully reproduced, like less easy recognition of hidden pockets and the emergence of many unrelated antibody sequences binding an antigen with high affinity. AM was correctly simulated at the scale of a complex 3D GC reaction. Realistic dynamics of individual GC cell numbers and affinity could be obtained. A substantial variation could be observed between simulated GCs, ranging from monoclonal bursts to polyclonal co-existence of clones, which is in line with the stochastic nature of GCs [32]. Each GC produced a different antibody sequence with high affinity, and some GCs performed better than others. The mutation path to high-affinity cells was not stringent and allowed for intermediate reductions of affinity. Along the path to high-affinity, typical affinity jumps of 2- to 10-fold and up to 74-fold were observed, which is associated with key-mutations in real GCs [36]. These results together provide evidence that the structure-related affinity landscape is sufficiently complex for studying structural implications of antibody development.

Thanks to new methods for enumerating and clustering possible peptide backbone structures [37], it will become possible to adapt the present algorithm to more realistic space conformations by discarding improper angles using rotamer libraries on finer lattices [38], and using approximate scoring functions for receptor-ligand affinities, as for protein docking. The use of real Amino Acids will allow to include out of frame and SHM hotspots into nucleotide sequences.

Several studies have used mutation trees of B-cell repertoires as readout of selection signatures [39]. The robustness of those methods may be improved by generating affinity matured repertoire benchmarks [40] with structurally related antigens and proposing the appropriate size and time-points of sequencing datasets to be analyzed [41].

Vaccine development is hindered by the hierarchical immunodominance of some epitopes over others, on the same antigen [42]. Some epitopes may be accessible and easily targetable, while others are hidden in pockets [43]. By reflecting accessible, shielded and hidden epitopes, the structural affinity model is suitable to study causes of immunodominance.

Immunodominance can also evolve over time, as observed during HIV vaccination [44]. Antibody feedback has been proposed to contribute to hiding immunodominant epitopes [18]. Our model can easily be extended to account for antibody feedback: The B-cell antibody sequence and an earlier produced antibody can compete to bind to the full antigen using mass action kinetics at equilibrium provided their target structures overlap, although such antagonism might be more complex [45]. In this setting, time-shifting of immunodominance will emerge naturally.

Vaccines against highly mutating pathogens like the hepatitis C virus, Dengue fever, influenza or HIV need to raise antibodies against many strains at the same time, ultimately towards the holy grail of broad neutralization. For instance, HIV envelope accessible regions are highly variable, while the conserved core pocket is harder to target [43]. The accessible immunodominant region diverts the immune system into recognizing regions that can escape easily without impact on viral function. Broad neutralization is induced by antibodies recognizing the conserved region like the CD4 binding site of HIV Env protein or influenza Haemaglutinin, thus binding most strains (conserved cross-recognition). But also cross-reactivity of antibodies that recognizes slightly mutated regions (promiscuous cross-reactivity) [46] or completely unrelated epitopes (poly-reactivity) [47] on various strains have the potential of broad neutralization. It is obvious that the best vaccination strategy depends on the antigen structure and the associated kind of cross-reactivity.

Simulations have the potential to inform the best vaccination strategy. Earlier works have simulated cross-recognition of conserved region with abstract affinities with a penalty on affinity for recognition of a shielding pocket [22]. Here, we have shown that GC simulations can be performed with antigens of variable and controllable structure and mutation landscape. This enables predictions of vaccine efficacy with cocktails of antibodies with any structural or mutation relationship, not only in the case of conserved recognition.

Further, by simulating waves of mutant viruses over a long period of time, it has been proposed [48] that the breadth of recognition by Tfh cells in GCs can determine the emergence of broadly neutralizing antibodies. More generally, the structural affinity model could be used to simulate the co-evolution of virus protein and GC B- and T-cells, as was performed at the repertoire level [21]. Altogether, the presented model paves the ground for computer simulations of AM and conditions that favor the development of bnAbs, thus, providing an advance in the field of *in silico* design of antibody-related immunotherapies.

## Methods

### 3D-lattice representation of proteins

Proteins structures are represented on a 3D Euclidean grid inspired from previous single-protein lattice models [24, 49]. Successive AAs occupy neighboring positions in the grid (Figure 1A). Only a single AA can occupy the same position. Starting from a grid point, a protein structure is represented as a sequence of moves on the grid, namely *straight* (S), *up* (U), *down* (D), *left* (L) or *right* (R). The first move can also be backwards (B). From predefined observer coordinates, each move is made relative to the previous observer coordinates and turns the observer in a new direction, except for a straight move (Figure 5A,B).

**Figure 5:**
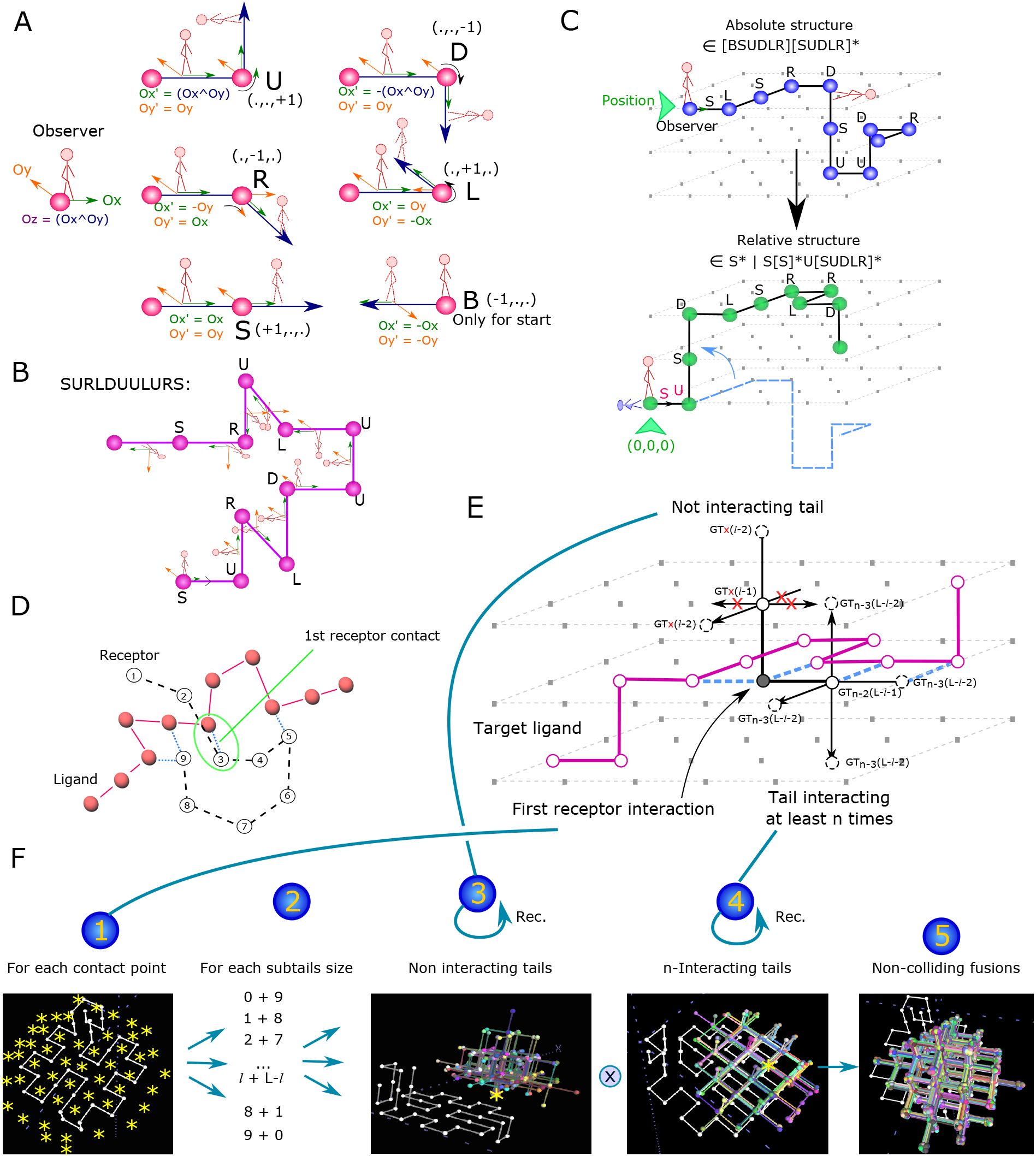
Representation of proteins and enumeration of receptor foldings. **A.** A protein structure is described as a list of ‘moves’ (see Figure 1B), that are relative to the observer coordinates 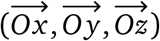. For each move, the new coordinates are given 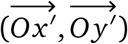 and 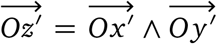. **B.** Example of the sequence ‘SURLDUULURS’ and its structure. **C.** Proteins can be represented as ‘Absolute’, with a starting position in space, or as ‘Relative’ after translation and rotation to start with S and followed by U, thereby representing all structures identical by rotation/translation in the same format. **D.** To enumerate all possible receptors binding a ligand, we define a ‘first position of contact’ to the ligand. Then, one of the two receptor tails does not interact with the ligand. **E.** Recursive rules for enumerating all possible receptors with a specific ‘first contact position’. All the possible structures for the two tails are enumerated separately, and each combination of structures from both tails is tried as a fusion to get a full, non self-colliding, receptor. The function Generate Tail (GT) of length *l* from a position *P* without contacts (red ‘x’) calls itself from the neighboring positions with length *l* − 1; similarly, the function GT of length *L − 1* with at least *k* contacts calls itself from neighboring positions with length *L* − *l* − 1 and *k* or fewer minimum contacts, depending on the number of contacts gained. **F.** Graphical illustration of the enumeration of all receptors described in E. For each possible first contact position, for each possible length of each tail with correct sum *L*, the possible interacting and non-interacting tails are enumerated from this position, concatenated and kept if they didn’t collide.

For each structure, a unique ‘absolute structure’ is defined by (1) a starting position and (2) a string built on the alphabet [SUDLRB], where B can only be the first letter. The ‘relative structure’ is defined as the group of absolute structures identical by translation or symmetry (Figure 5C). This is achieved by forcing the first move to be ‘S’ and the first turn to be ‘U’, thereby removing the ‘B’. Relative structures follow the regular expression: ‘S* | S[S]*U[SUDLR]*’ and do not need a start position on the grid.

The use of relative structures has several advantages. It allows to enumerate and manipulate possible structures by generation of random strings according to the regular expression. The representation is compact (an alphabet of five letters) and collisions (two AAs at the same position) among three successive AAs are avoided by removal of ‘B’. Rotations around a covalent bond are performed by changing a single letter, inducing a turn of the observer coordinates and propagating changes of the turn names according to the new coordinates, but without computing the new spatial positions of the AAs. It is also easy to fuse two proteins into a longer one: All possible structures resulting from the fusion, according to different rotations, can be found by fusing the sequences and performing turns of the observer coordinates. It has to be noted that the same protein structure can be described from both ends, and the sequence of a relative structure can be easily reversed to describe it by starting from the other end.

### Binding and total energies between two proteins

For two proteins (as in Figure 1), an interaction is defined by a pair of two non-consecutive AAs occupying neighbor positions. Interactions can be within a protein (folding) or between two proteins (binding). To distinguish the notions of ‘empty’ structure and protein (that is a structure with AAs), we introduce the operator *P*(*R*, *S*) that represents the protein of structure *S* and AA sequence *R* (both sharing the same length). The binding energy *E*_bind_ between a receptor protein *P*(*R*, *S*) of length *L* and an antigen *P*(*G*, *K*) of length *L*_G_, is the strength of the interaction between the two proteins and is calculated as the sum of all interactions between them:

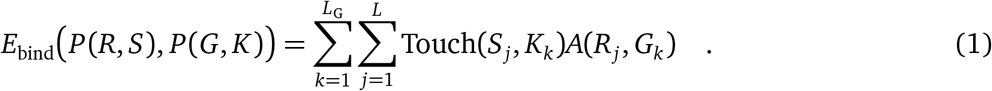

The *i*^th^ grid positions of a structure *S* is denoted by *S_i_*, irrespective of the type of AA at this position. The operator Touch(*S_j_*, *K_k_*) returns 1 if the residues *S_j_* and *K_k_* are non-covalent neighbors. *A*(*R_j_*, *G_k_*) is the interaction potential between the residues types given by [30]. The folding energy *E*_fold_ of a protein *P*(*R*, *S*) of length *L*, is the sum of intrinsic interactions between its own interacting AAs:

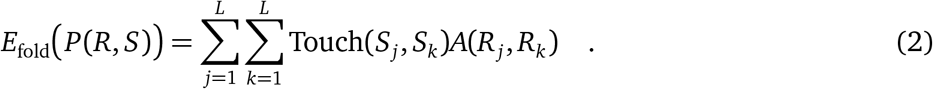

As we assume a static conformation of the antigen, we can define the ‘total energy’ *E*_tot_ of *S* around the antigen *P*(*G*, *K*) as the sum of its binding and total energies:

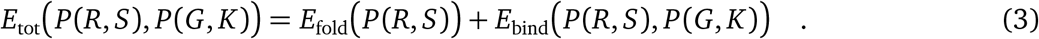

Energies are negative and calculated in *kT* units. Strong binding implies low energies.

### Combinatorial enumeration of all possible foldings

From a predefined ligand, we explicitly enumerate all the possible 3D structures of the receptor of length *L* with at least *n*(≥ 1) interactions with the ligand. Starting from the first receptor residue, its first interaction point with the ligand is localized (Figure 5D). By definition, further interactions with the ligand have to be behind this point.

From a particular empty grid point *X* that is neighbor of a ligand residue, it is possible to recursively enumerate all receptor structures with *X* the first contact point to the ligand. All structures without interaction until *X* and all structures that may interact with the ligand behind position *X* are enumerated. Next, each pair of starting and finishing structures are checked for collisions with each other and for the right total length *L* (Figure 5E).

The recursive algorithm ‘GenerateTails’ (Algorithm 1) shows how to get a starting or finishing structure, and the algorithm ‘GenerateReceptors’ (Algorithm 2) shows how to enumerate all possible receptor structures around a ligand, with a minimum number of interactions between receptor and ligand. In order to improve efficiency, the GenerateTails function will be called multiple times from the same point, and the result of each call is stored in memory, avoiding excessive recomputing and explaining why memory usage reaches a few GB (Figure 1G).

Note that each starting point *X* describes a set of mutually exclusive structures if we assume oriented receptor structures (first to last residue). Indeed, every structure, if not oriented, would be enumerated twice, once from each end. Algorithm 2 is still valid in the particular case of *n* = 1 because the structures will be enumerated twice in lines 18-23.

Finally, in order to minimize the computational time, a list of structures can be compressed into a list of binding pairs of positions on the receptor and AAs on the ligand. Some structures produce the same list of binding pairs. By keeping track of which structures have which binding pairs in a dictionary, only the binding pairs need to be stored and evaluated for further exploration of the binding energies of concrete receptor sequences. This step leads to a three- to four-fold increase in computational speed.

**Algorithm 1.**
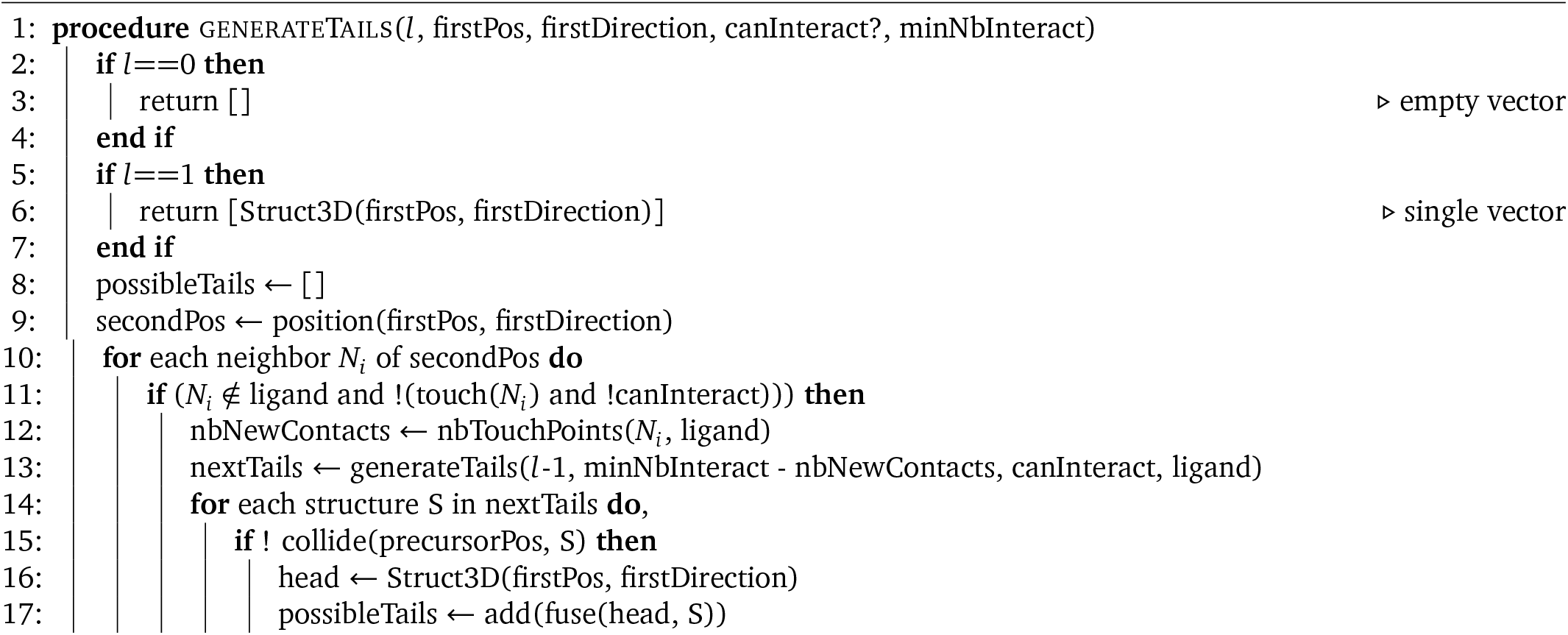
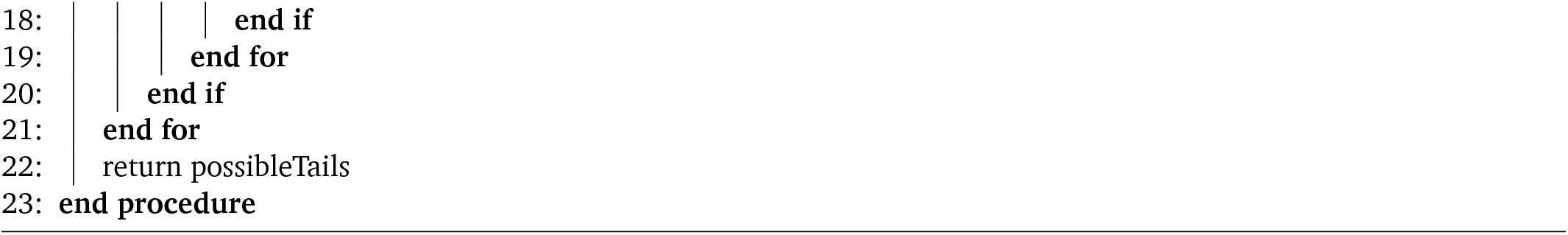
Function that generates the list of all possible structures of length l that start from a specific point and direction, and interact at least *n* times with the protein or that do not touch the ligand.

**Algorithm 2.**
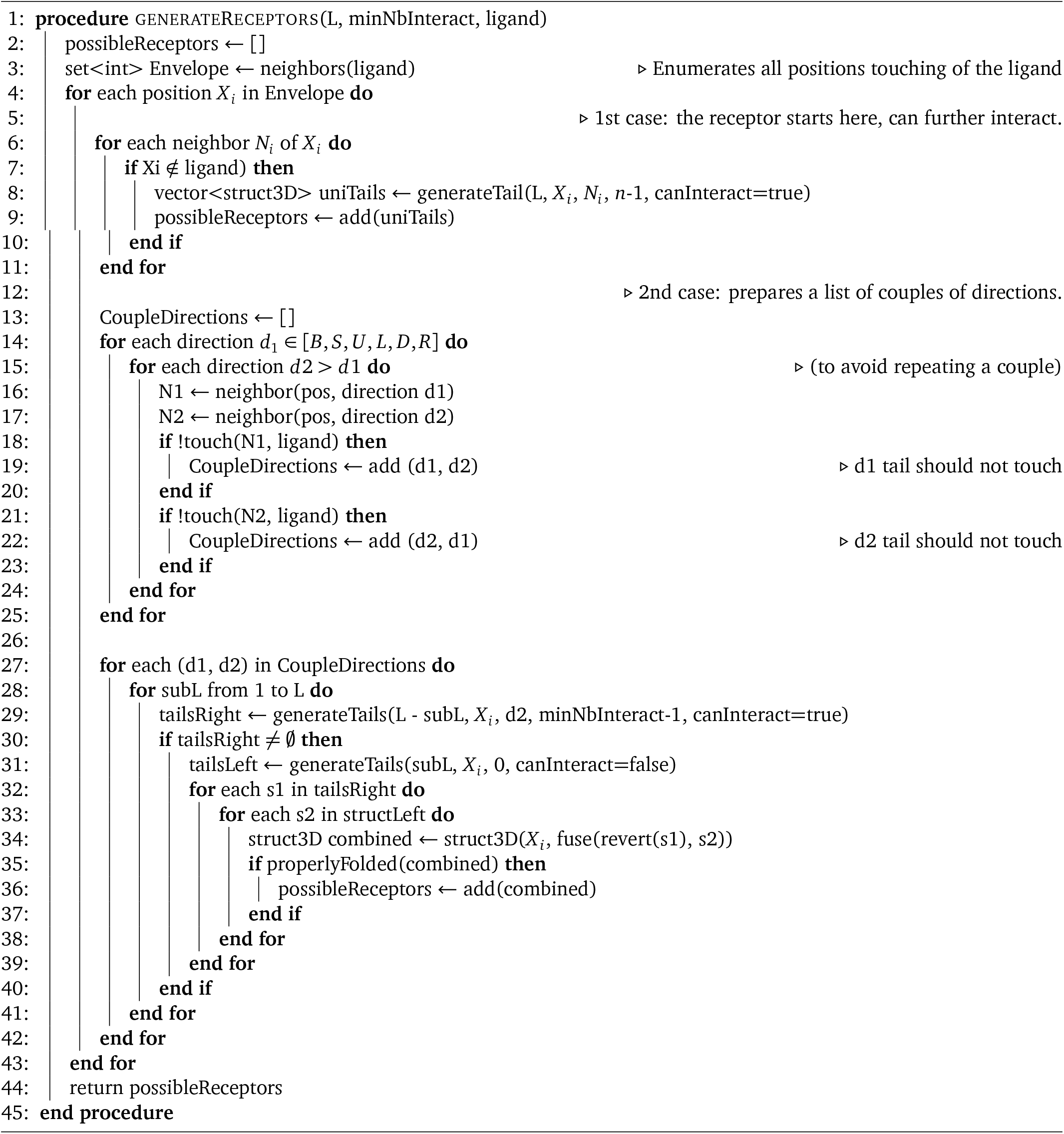
Main function to enumerate all receptors structures of length *L* that interact at least *n* times with the ligand.

### The best binding energy of a receptor sequence to the ligand

Given the pre-computed list of possible 3D receptor structures with at least *n* interactions to the pre-defined ligand, the binding energy and total energy between the two AA sequences is calculated for each structure with the receptor AA sequence. The ‘best binding energy’ *E*_best_ is the average binding energy of all optimal structures in terms of ‘total energy’ *E*_tot_ (around a small *ϵ* = 0.001 to account for rounding errors), with

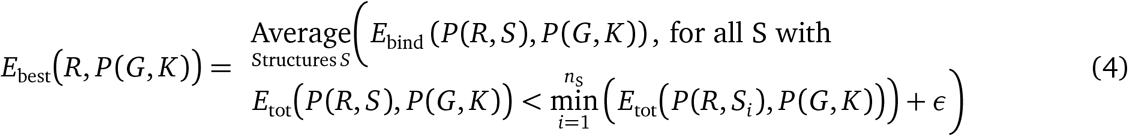

Self-folding can contribute to stabilize a structure and therefore the best structure is not necessarily the one binding with highest strength, but with lowest total energy. *R* is the receptor sequence, *G* the antigen sequence, *K* the antigen structure, *S* are the possible enumerated receptor structures, and *n*_S_ is the number of these structures. If a receptor sequence folds on itself with better energy than the total energy around the ligand, no binding is assumed and the energy is *NAN* (Not A Number).

Alternatively, a ‘statistical energy’ can be computed by applying a Boltzmann weight to each protein structure *P*(*R*, *S_i_*) according to its total energy:

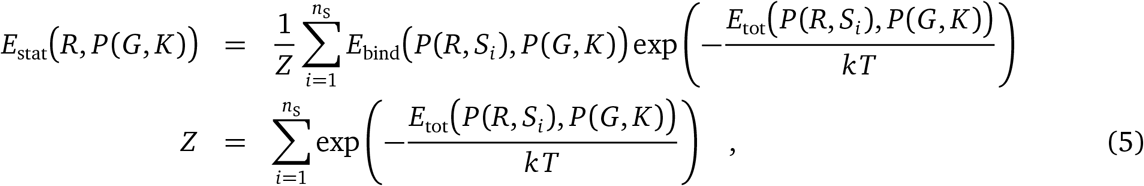

with *kT* the Boltzmann coefficient and *S_i_* the *i*^th^ structure. *Z* is the sum of weights among all possible structures, and is used as normalization coefficient. Many structures could co-exist with good but not optimal total energy, and this may not be described well by the set of optimal structures alone in *E*_best_. In theory, the Boltzmann distribution should be distributed according to all possible receptor structures, rather than only with respect to a particular minimum number of interactions *n*, even including those that do not interact with the ligand (self-foldings). Structures with *n* < 4 had no effect on Z (not shown) because these come with a small weight. We also computed all the possible self-foldings for different receptor sizes *L*, and found that the influence of *n* on *Z* was negligible (not shown) because the number of possible self-folding structures was small compared to large number of foldings around the ligand. This justifies to discard self-foldings and choose *n* = 4. Higher values of *n* may be used to increase speed, as shown in Figure 1F, provided that there is still no impact on *Z*.

### Transforming energy into affinity and binding probability

The best binding energy between receptor and ligand was calculated for short receptor structures. The real antibody–antigen affinity needs to include the contribution of the full antibody, with two binding regions and the scaffold. Further, inside a GC, the affinity of a BCR to the antigen translates into capture and internalization of the antigen, a process that depends on many other factors. Therefore, an affinity cannot be drawn easily. We define an empirical ‘re-scaled affinity’ a

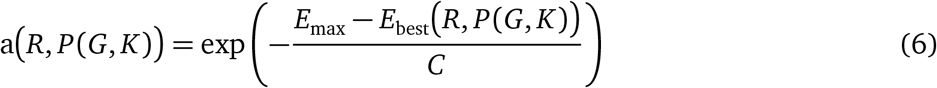

of a receptor AA sequence *R*, to represent the probability of binding the antigen. *E*_max_ is the binding energy, at which the antigen capture probability becomes 1. The binding energy was re-scaled with a coefficient *C* because the best structures were dominant in the affinity and GC simulations did not develop because of low antigen binding. In Figures 3 and 4 the affinity of L2, L5, and L6 and receptors with *L* = 9 were calculated with: *E*_max_ = −100 and *C* = 2.8. *E*_max_ and *C* have to be adapted for different receptor length *L*.

### GC simulations

GC simulations were performed on an agent-based model programmed in C++ in the lab, for which the algorithm was described in [9, 50]. The model explicitly simulates the movement and encounters of B- and T-cells, capture of antigen by B-cells, T-cell help, proliferation, recirculation, death and exit from the GC. The model settings with constant inflow of founder cells and a single B-cell - T-cell interaction for B-cell selection were used [51]. Parameters values are detailed in [50].

The structural affinity model replaced the shape space affinity model in this framework. A mutation is a random change in the AA located at a randomly picked position of the BCR. Selection parameters had to be adapted: Antigen uptake is limited in the model by a ‘refractory time’ between two attempts of antigen capture. This time was reduced to 3.6 seconds. The simulation time-step had to be lowered to 0.001 hours accordingly.

In the model, the individual number of B-cell divisions upon T-cell selection is derived from a Hill-function depending on the amount of captured antigen. We observed that the appearance of high affinity B-cells happened slower with the structural space compared to the shape space. We needed to lower the Hill-coefficient from 2.0 to 1.4. This value was also supported in [52]. With a higher slope, cells with intermediate affinity did not expand (not shown).

Each founder B-cell entering the GC carries a randomly picked BCR of length *L* = 9 AAs and with a minimum affinity of 0.0001 to at least one epitope, potentially allowing for an increase in affinity by a factor of 10,000 in a GC reaction. If we lower this ‘entry’ threshold further, the simulated GCs collapse and the mutations do not reach reasonable affinities (data not shown).

In the model, Follicular Dendritic Cells (FDC) occupy a set of positions in space, and display a certain amount of epitopes at each of position to the B-cells. When using multiple epitopes, the same total amount of epitopes in the simulation is used as for single epitope simulations, meaning two epitopes are used each with half-amount, etc. Each FDC position was initially filled with an equal amount of each epitope. At each position, B-cells can access all of the epitopes simultaneously. The antigen capture probability was determined by the highest affinity to any of the epitopes at this position, and this highest affinity epitope is removed by one epitope unit at this position. Other epitopes are not captured.

Computation times are given for simulations on a single Intel Xeon CPU core (model E5-2690 v3 at 2.6 GHz). The antigen sequences used for 4 are given in Suppl. Table S1.

### Graph analysis

Each founder sequence was assigned a unique ID. For every mutation, a new unique ID is assigned to the mutated sequence, even if this sequence already exists in another cell. The mutation history network of one GC is created with sequence IDs as nodes and mutations as edges. It is a forest, where each founder dynasty is a tree. The network was analyzed with Cytoscape (Figure 3). To represent the network, the default Prefuse Force directed layout was used, which showed a cluster for each founder in a convenient way. For Figures 3C and G, a cluster was manually selected, the nodes with an affinity of more than 0.5 were selected and their parents were included up to the founder sequence. The Prefuse Force directed layout was used to represent the graph. For Figure 3F, the tree in Figure 3E was selected, the nodes with identical sequences were merged and the network was shown again. The sequence logos (consensus) of Figure 3B were generated using Skylign (skylign.org).

## Acknowledgements

This work was supported by the Human Frontier Science Program (RGP0033/ 2015), and a PhD fellowship granted by École Normale Supérieure de Lyon. We thank Victor Greiff, Rahmad Akbar and Gang Zhao for fruitful discussions and suggestions, and Megan Foster (LeafItToMe) for critical reading of the manuscript.

## Author contributions

PR and MMH designed the study, MMH programmed the Germinal Center model, PR programmed the structural affinities, MMH and PR analyzed the results and wrote the manuscript.

## Supplementary material

**Figure S1:**
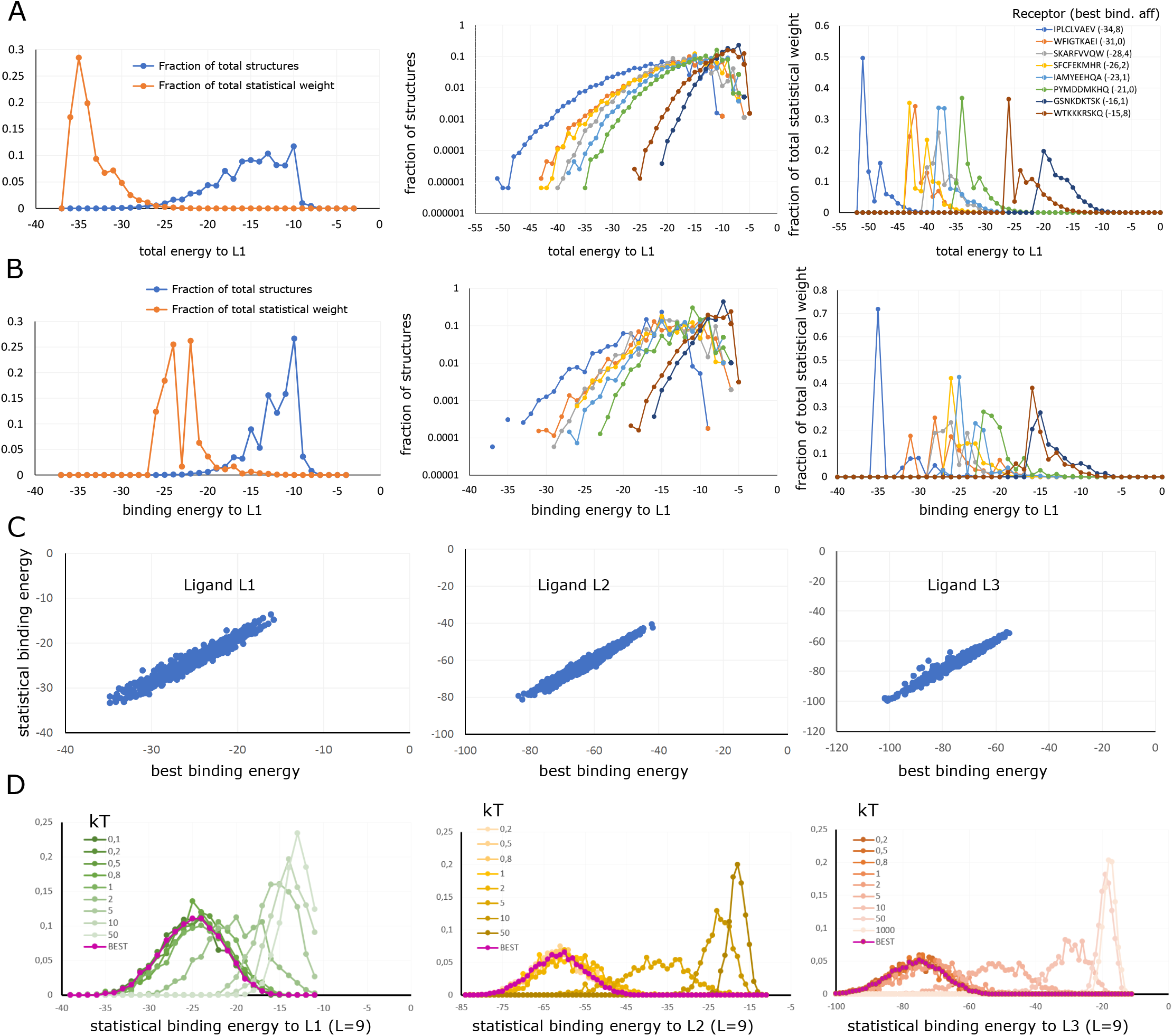
The best binding energy is a fair approximation of the statistical binding energy. **A-B.** Description of the energies of the ensemble of structures around ligand L1, for particular receptor sequences. **A.** Left: Distribution of total energies *E*_tot_ and statistical weight for possible structures of one receptor sequence. Most structures have higher (non-favorable) total energy, but only the ones with lowest energy have a significant statistical weight. Middle and right: Distribution of the total energy *E*_tot_ of the possible structures for selected receptor sequences with different best binding energies, as fraction of the total statistical weight *Z* (middle) or as fraction of the total number of structures (right). **B.** Same analysis with binding energies *E*_bind_ instead of total energies. **C.** Correlation between the best and statistical binding energy for randomly selected receptor sequences of size 9 against ligands L1, L2 and L3. **D.** Effect of temperature on the distribution of statistical binding energies of randomly picked receptors sequences of size 9 with each ligand. Temperature values around *kT* = 1 have no effect on the distributions, meaning that a frozen state is reached, where optimal structures dominate.

**Table S2:**
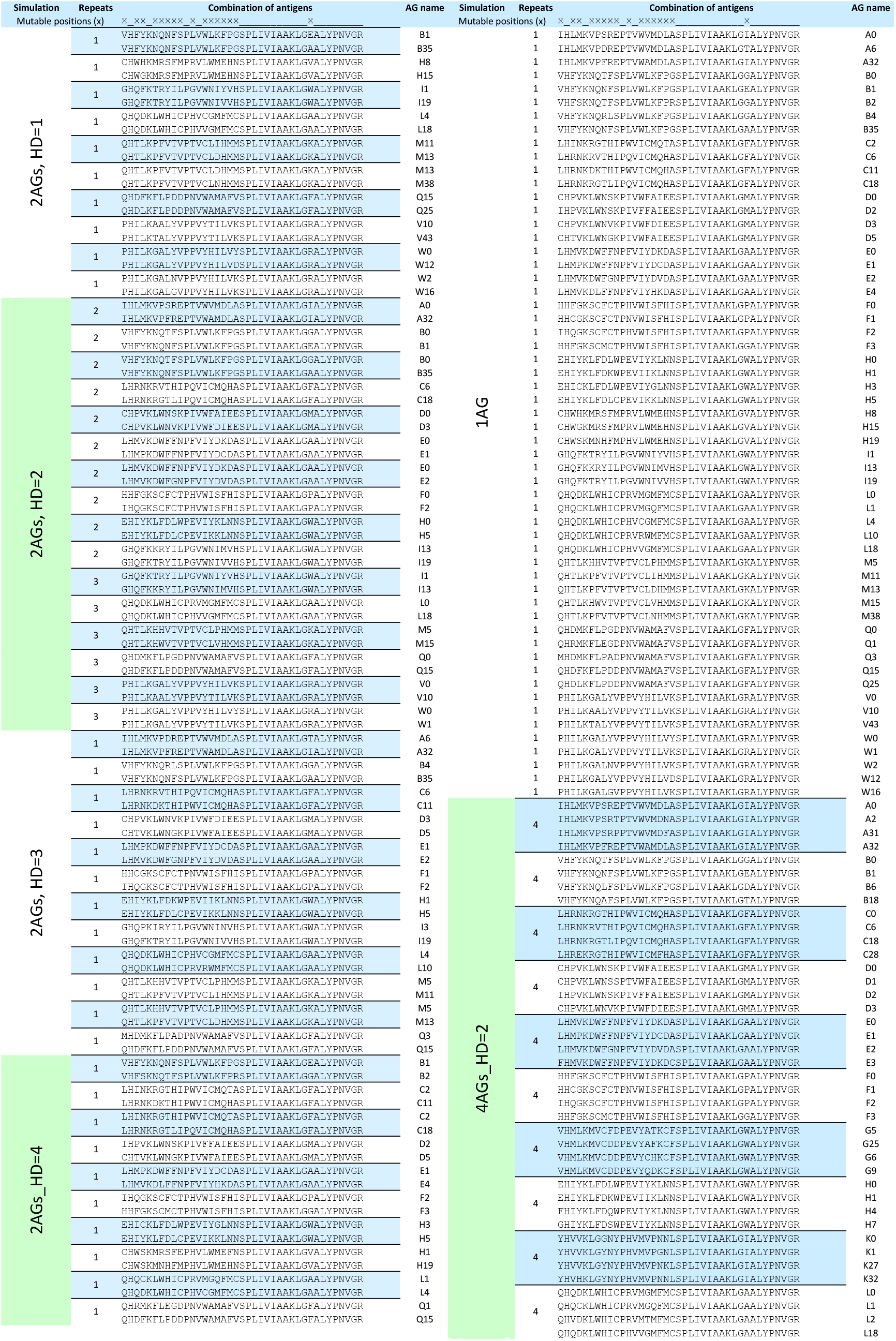
Antigen sequences used for GC simulations. For each group of simulations (left column), the set of antigen sequences used on structure L6 is given, together with the number of replicates of each GC simulation. Briefly, antigen AA sequences that achieve an average best binding among 200 random receptors between the top 1% and 15% best values were selected, to avoid chaotic GC dynamics if the antigen has too low or too high binding, and sequences with 1 to 5 point mutations within this range were kept to be used as combination of antigens with different Hamming Distances (HD).

